# Parasites Mediate Condition-Dependent Sexual Selection for Local Adaptation in a Natural Insect Population

**DOI:** 10.1101/563874

**Authors:** Miguel Gómez-Llano, Aaditya Narasimhan, Erik I. Svensson

**Author notes:** Data Accessibility Statement: Data will be accessible in a public repository upon acceptance.

## Abstract

Condition-dependent sexual selection has been suggested to reduce mutation load, and sexual selection might also accelerate local adaptation and promote evolutionary rescue through several ecological and genetic mechanisms. Research on condition-dependent sexual selection has mainly been performed in laboratory settings, while data from natural populations are lacking. One ecological factor that can cause condition-dependent sexual selection is parasitism. Here, we quantified ectoparasite load (*Arrenurus* water mites) in a natural population of the common bluetail damselfly (*Ischnura elegans*) over 15 years. We estimated parasite-mediated sexual selection in both sexes and investigated how parasite resistance and tolerance changed over time and influenced population density. Parasites reduced mating success in both sexes, but the effects were stronger in males than in females. The male fitness advantage of carrying fewer parasites was higher under experimental low-density conditions than under high-density field conditions, suggesting that male-male competition could reduce parasite-mediated sexual selection. We further show that population density declined during the study period, while parasite resistance and male fitness tolerance (fecundity) increased, suggestive of increasing local adaptation against parasites and ongoing evolutionary rescue. We suggest that condition-dependent sexual selection can facilitate population persistence and promote evolutionary rescue by increasing local adaptation against parasites.

## INTRODUCTION

Theory suggests that sexual selection can promote local adaptation [1,2] and can facilitate evolutionary rescue [3], for instance, by purging the genome from deleterious mutations [4]. The negative fitness effect of all deleterious mutations carried by a population, or the mutation load, can have detrimental consequences, potentially leading to population extinction [5,6]. Sexual selection can reduce mutation load if mutations that are deleterious to non-sexual fitness also reduce mating success [4,7–10]. Sexual selection against deleterious mutations could facilitate evolutionary rescue by promoting local adaptation [4,8,11–15]. As mean population fitness is more closely linked to female than to male relative fitness, sexual selection can reduce mutation load without substantial demographic costs and without affecting population productivity [16]. If the strength of selection is stronger in males than in females and provided that females mate with other males, then females can benefit from sexual selection on males that carry most of the costs of deleterious mutations, while the mean population fitness remains unaffected [4,17,18].

Previous theory and some empirical work, primarily from laboratory settings of model organisms (see above), have focused on the effects of sexual selection in reducing the mutation load and the links to local adaptation. However, it is worth underscoring here that the role of sexual selection can be much broader, as it can also influence population viability, local adaptation and extinction risk through more ecological pathways [19–21], in addition to the above-mentioned population genetic mechanisms. For instance, the rapidly growing interest in eco-evolutionary dynamics highlights the feedbacks between ecological processes and evolutionary change [22]. Eco-evolutionary dynamics has only recently become extended to the fields of sexual selection and sexual conflict [19,21]. A small but growing body of literature shows that sexual selection and sexual conflict can indeed have ecological consequences for female fecundity and population fitness [16,23–25], community structure [26,27] and for parasite loads [21]. These ecological consequences of sexual selection and sexual conflict have only recently begun to be explored more systematically in terms of the consequences for local adaptation and evolutionary rescue [20,28]. Thus, there are good theoretical and empirical reasons to expect cascading ecological consequences and eco-evolutionary feedbacks from sexual selection and sexual conflict to ecology, and such effects can have community-wide effects [19,21].

Sexual selection studies have often focused on secondary sexual characters, such as signalling traits, although overall condition is likely to contribute more to individual mating success [4,9]. As mating success require males to succeed in several energetically costly behaviours (e.g., searching for females, male-male competition, courtship, female coercion, etc. [29]), male mating success is likely to be highly condition-dependent [30]. There is ample evidence that the expression of sexually selected traits is highly dependent on overall condition [29,30], which is a strong determinant of individual fitness, and is therefore likely affected by most loci across the genome and are presumably also target of directional selection. Because of this directional selection pressure towards higher condition, novel mutations in a population are more likely to reduce condition than to increase it [30].

Many behaviours influencing mating success (e.g., mate searching, male-male competition) can vary in intensity with density, affecting the strength of sexual selection [31–33]. For example, in high-density populations mate-searching costs might be reduced, leading to weaker sexual selection, but competition could also become more intense which would increase the strength of sexual selection. Few studies have empirically investigated density-dependent effects of sexual selection, and results are contentious. Notably, a study by Sharp and Agrawal [8] on laboratory bred *Drosophila melanogaster* populations found no difference in strength of sexual selection between high and low density conditions. Another laboratory study on *D. melanogaster* investigated the effect of searching costs in sexual selection (searching costs are expected to be higher at low densities). The authors found increased strength of sexual selection when searching costs where higher [9]. In a study with fungus beetles (*Bolitotherus cornutus*) in six different natural populations, Conner [34] found stronger sexual selection for long-horned males in low densities, where usually only one long-horned male was found. However, in high densities, intense competition with other long-horned males reduced male mating success [34]. Density-dependent sexual selection, especially in natural populations, is clearly an area that requires more research before any firm conclusions can be made.

Parasite load has detrimental consequences on fitness, reducing hosts’ condition, which can often also affect mating success [35,36]. For instance, field studies on red grouse (*Lagopus lagopus*) revealed a negative effect of parasite load on body condition and resulted in condition-dependent expression of secondary sexual characters [37]. In laboratory bred populations of *D. melanogaster*, experimental infection with the bacterium *Pseudomonas aeruginosa* revealed sex differences in parasite-mediated sexual selection [38]. The aforementioned studies suggest that parasite load can reduce condition, and as a result, influence sexual selection.

Pond damselflies (family Coenagrionidae) to which *I. elegans* belong, are characterised by strong sexual selection, arising from intense male-male scramble competition over access to females and associated male mating harassment [31,32,39,40]. Mating success is therefore highly likely to be condition-dependent, given intense male-male scramble competition in damselflies [41,42]. Natural populations of pond damselflies are often infected by ectoparasitic water mites of the genus *Arrenurus* [43–45]. High parasite load can affect agility and flight performance, which is necessary for mate search and competition for mates [44]. In a previous study in the *I. elegans* and the *Arrenurus sp.* system, we found a negative effect of ectoparasites on female fecundity [46]. However, the effects of parasite load on male mating success and sex differences in the strength of sexual selection against parasitism have not been investigated before.

Here, we used long-term observational data from a natural *I. elegans* population that has been monitored for 15 years to investigate whether male and female parasite load affects either precopulatory (mating success) or postcopulatory (fecundity) sexual selection. We present a comprehensive, multi-generation field study showing evidence of condition-dependent sexual selection caused by parasite load and its effects on population density, local adaptation, parasite resistance and parasite tolerance in both sexes. We quantified the strength of selection against parasite load using selection differentials and compared the strength of pre- and postcopulatory sexual selection in the two sexes. We complemented these analyses performing mate choice experiments under low-density conditions to confirm the role of parasites mediating sexual selection on males. Finally, we looked for signatures of increased local adaptation through temporal changes across years in the parasite load and the effects on fecundity in this population and in the link to changes in population density. Based on our results in this study, we suggest that parasite-mediated and condition-dependent sexual selection can promote local adaptation for parasite resistance and tolerance, and thereby contribute to population persistence and decrease extinction risk.

## METHODS

### Study organism

The common bluetail damselfly (*I. elegans*) is a small insect species distributed throughout Europe, with its northernmost range in central Sweden [39,47]. The generation time of *I. elegans* is one year in the northern part of its range in Europe, with adults emerging between late May and early August, while the larvae overwinter in an aquatic stage [39]. During the aquatic stage, water mites of the genus *Arrenurus* attach to the larvae and remain in a phoretic stage until adults emerge, when they attach to the cuticle on the ventral side and pierce it to feed on the host body fluids [43–45]. Water mites are easily detected visually, and have been reported to affect agility and flight performance in other species of *Coenagrion* and *Enallagma* damselflies [44]. In *I. elegans* specifically, these mites have been demonstrated to have negative effects on female fecundity [46].

Mating and reproduction in *I. elegans* consists of several behavioural steps. First, males search for females, which they attempt to clasp by the prothorax using structures known as claspers, located at the terminal abdominal segment [39]. Intense competition between males to find and clasp females is a typical feature of the mating system of *I. elegans* and most other pond damselflies that have been studied in detail [39]. After a successful male clasping event, the male and the female form a tandem, and the female can then choose to bend her abdomen to reach the male genitalia and ensure mating and sperm transfer [39]. During copulation, male damselflies remove sperm from previous matings before transferring their own sperm to a female [48,49]. Finally, after mating, females oviposit with no male mate guarding on emergent vegetation in freshwater bodies [39].

### Data collection

Our observational data come from a long-term study of natural populations of *I. elegans* collected over the last 15 years (2003-2018) in Southern Sweden. The location used in this study is a large permanent freshwater pond located in Lomma (55.684942, 13.085414). We visited this population in Lomma on multiple occasions during the 15 reproductive seasons of *I. elegans* (early June to early August). During these visits, damselflies were caught using hand nets. We caught all *I. elegans* males and females we could during time-recorded capture sessions, and estimated population densities dividing the total catch with the catching effort (*i. e.*, individuals caught per minute). For each captured individual we identified the sex, mating status (single or in copula), and number of parasites. Females captured *in copula* were transported back to an indoor laboratory at the Department of Biology (Lund University) were we estimated fecundity by placing females in individual plastic cups with moisturised filter paper as an oviposition substrate [50]. Females were allowed to lay eggs for 72 hours, after which the eggs were counted. Fecundity was only measured of those females that were found mating in the wild, as we could quantify the parasite load in both parents. Mating success and parasite load was collected from 2003-2018, but fecundity estimation could not be collected during the period between 2010 and 2013.

### Experimental mating trials

We performed a series of mating trial experiments under low-density conditions (one female with two males) during the reproductive season (June and July) in 2018. We collected parasitized and non-parasitized males and females from the Lomma population. Captured individuals were then transported to Stensoffa Field Station (55.695145, 13.447076) in netted containers (“*Port a bug*” #T-11-232, 10.2 cm diameter and 22.9 cm height). All animals were separated by sex and kept in an average density of 10 individuals per container during the transport to the field station. We quantified the parasite load of each individual and marked the females randomly with fluorescent colour dust in the genital area, approximately on the last 2-3 segments of the abdomen. This technique allowed us to identify the males who successfully mated by looking at traces of colour dust in their genitalia under black light. This technique has previously been used to quantify mating success and mating rates in *I. elegans* and other damselfly species such as *Enallagma cyathigerum* and the genus *Calopteryx* [26,32,51]. We set up a series of cages (above mentioned “*Port a bug*” netting containers) for mating trials, which consisted of one marked female, one parasitized male and one non-parasitized male. We added twigs and grasses to mimic natural vegetation and allow individuals to perch or rest, and a plastic cup with a moisturised filter paper to prevent desiccation. After 24 hours, we inspected each individual, and classified each male as mated or unmated, depending on whether traces of the fluorescent dust carried by the female was observed on the male’s genital area.

### Statistical analysis

All the statistical analyses were performed using Bayesian generalised mixed models with Markov Chain Monte Carlo estimation with the package “MCMCglmm” [52] in R [53]. We used an uninformative prior, and performed 100,000 iterations, with a burn-in of 10,000 and 10 thinning interval. We evaluated convergence by plotting the chains and assessing if they mixed properly.

Precopulatory sexual selection was analysed with a binomial distribution using mating success (1 = mated, 0 = unmated) as a response variable, parasite load, sex and their interaction as fixed effects. To investigate density-dependent effects and compare the results from the experimental mating trials with sexual selection in the field, we used a model with binomial distribution using male mating success as response variable, parasite load, type of data (field or experimental mating trial), and their interaction were treated as fixed effects. To investigate postcopulatory sexual selection we used a subset of the data including only those females that were found mating to quantify how parasite load in the female and the male of the copulating pair influenced the female’s fecundity (number of eggs laid in the laboratory). We used a model with Poisson distribution, with number of eggs as response variable, parasite load in the female and the partner were used as fixed effects.

To estimate the linear selection differential [54] on parasite load, we regressed the relative fitness (mating success and fecundity) and standardized parasite load in males and females. As significance levels can be unreliable when using binomially distributed variables, such as mating success [54], we used binomial distribution to estimate selection differentials of mating success [31]. In this study, we are interested in the strength of selection differentials rather than their significance. Selection estimates can be very useful in meta-analysis that incorporate different taxa, traits, time and selection pressures into sexual selection (see [55–60] for examples). In all models specified above, year was included as random effect to account for statistical non-independence.

To investigate the consequences of sexual selection in local adaptation we performed two different models to analyse parasite resistance and tolerance separately from 2003 to 2018. Parasite resistance was analysed using a binomial distribution with parasite prevalence (1 = parasitized, 0 = non-parasitized) as response variable, year, sex and their interaction as fixed factors. Parasite tolerance was analysed as the reduced negative effect of parasite prevalence, seen through fecundity. We used egg count as response variable assuming a Poisson distribution. Year, parasite prevalence in females and males, as well as the two-way interactions (year and parasite prevalence in females, and year and parasite prevalence in males) were included as fixed factors. Finally, to assess population density changes through time, we regressed population density (number of individuals observed divided by the length of the sampling period for each visit) against year of study, assuming a normal distribution.

## RESULTS

We collected a total of 4387 individuals, 2655 males (1006 parasitized, ∼38%) and 1732 females (817 parasitized, ∼47%). Parasitism was less frequent in males than in females. We estimated fecundity of 554 females, which were captured *in copula* in the field. As we aimed to analyse the effect of male and female parasite load in fecundity, only females captured together with their partner and in which we were able to estimate parasite load of both parents were used for fecundity analysis. The mating trial experiments were performed with 70 females and 140 males, 70 of which were parasitized and 70 were non-parasitized.

### Sex differences in the effects of parasite load on mating success

Precopulatory sexual selection was estimated as male and female mating success (a cross-sectional measure, based on the proportion of copulating individuals in the field). We found a significant negative effect of parasite load on mating success (pMCMC < 0.001). We also found a significant effect of sex, where the average mating success of non-parasitized males was significantly lower than of non-parasitized females (pMCMC < 0.001), reflecting that a higher fraction of females were found mating near the water bodies where sampling took place. Interestingly, the interaction between parasite load and sex (i.e., testing for a sex-difference in slopes) was also significant (pMCMC = 0.024), revealing that the costs of parasite load to mating success is larger in males than in females (Fig 1A, Supplementary Table 1).

**Figure 1.**
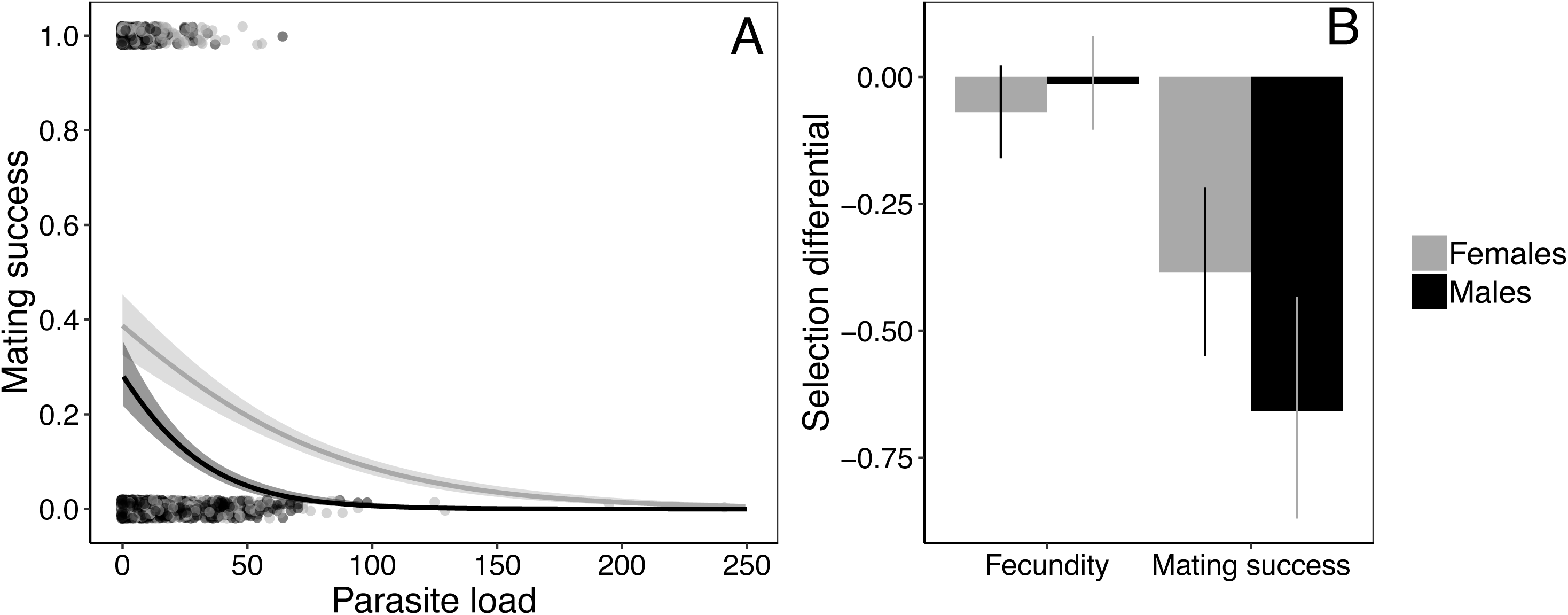
Parasite load reduced both male and female mating success under field conditions of male-male scramble competition. Results show a more negative effect of parasites in males than in females (i.e., steeper slope of the relationship between mating success and parasite load)**(A)**. The selection differentials against parasite load for mating success were significant in males and in females, while the selection differentials for fecundity were not significant **(B)**. Models were performed assuming a binomial (for mating success) and Poisson (for fecundity) distribution. Error bars and shaded areas show 95% confidence intervals. Points show individual observations and the regression line show predicted values.

Selection differentials (standardized slopes) for mating success against parasite load differed significantly between males and females, being stronger in males than (pMCMC = 0.031; Fig. 1B), consistent with the steeper slope for males in males than in females (Fig. 1A). Postcopulatory sexual selection against parasitism was estimated from individual fecundity data of females found in copula and their mates. We did not find a significant effect of neither male (pMCMC = 0.82) nor female parasite load on fecundity (Fig. 1B; pMCMC = 0.19; Supplementary Table 2).

### Parasite loads and male mating success in the field and in experimental trials

We compared the effects of parasite load on male mating success under high-density conditions of scramble competition (field data) with the effects of parasitism on male mating success under low-density conditions (experimental trials with one female and two males). Overall, we found a significant negative effect of parasite load on male mating success (pMCMC < 0.001) and a significant difference between the experimental trials compared to the field, with a lower male mating success in the latter situation (pMCMC < 0.001). We also fond a significant interaction between parasite load and density (low-density experimental trials and high-density field data), with the negative effect of parasites being less pronounced in the observational data in the field than in the experimental trials (pMCMC = 0.02; Fig 2A, Supplementary Table 3).

**Figure 2.**
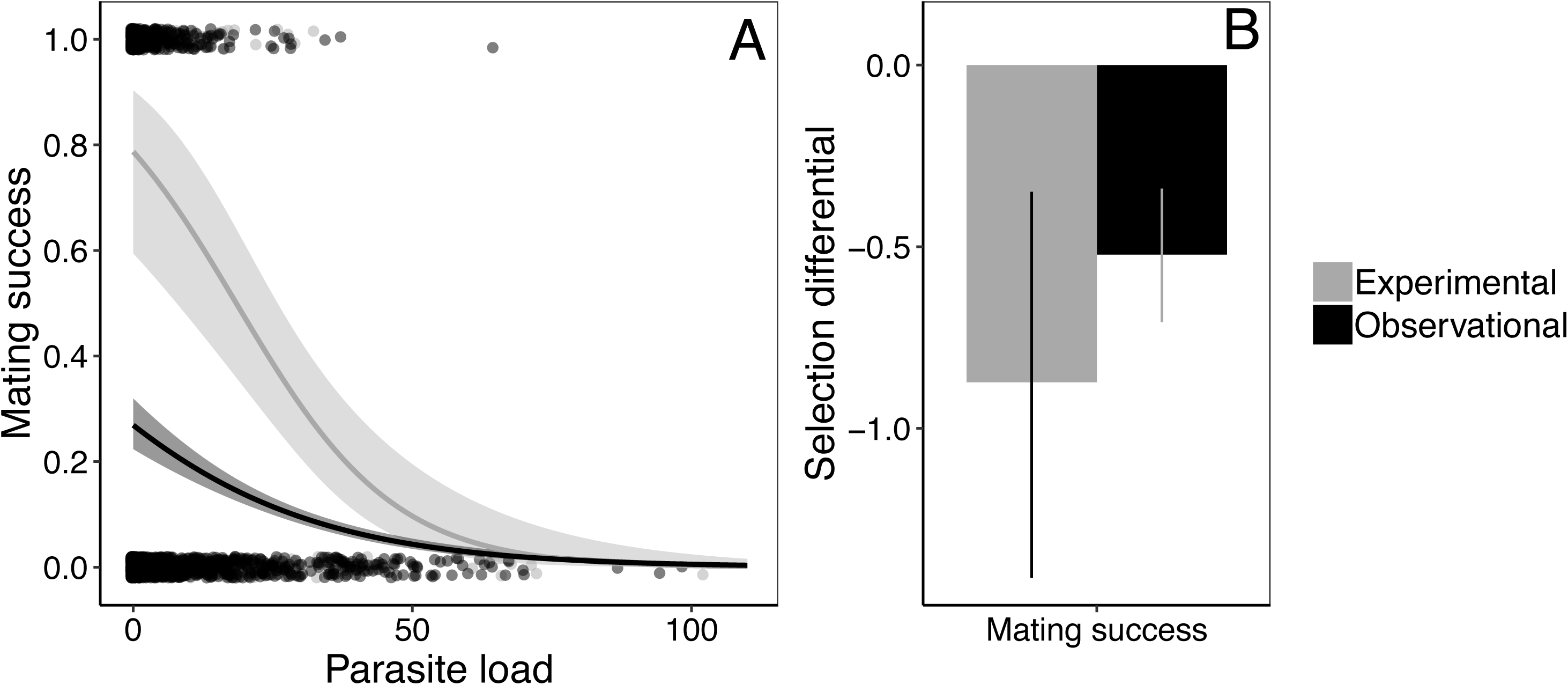
Mean mating success in non-parasitized males was higher (larger intercept) and the negative effect of parasite load more pronounced (significantly more negative slope) in the low-density experimental settings than in the observational data from the field. (**A**). Selection differentials of the effects of parasite load on mating success confirm that precopulatory sexual selection against parasitized males was stronger under low-density experimental conditions (one female with two males) than under field conditions with male-male scramble competition (**B**). We used binomial models in a Bayesian framework. Points show individual observations and the regression line show predicted values with 95% confidence intervals in shaded areas and error bars.

Finally, we compared the strength of selection against parasitism in these experimental low-density trials with selection estimates from the field. Indeed, precopulatory sexual selection against parasitized males through mating success was more than three times stronger in these low-density experimental trials (*s* = −1.8) than in the high-density field data (*s* = −0.57; Fig 2B, Table 2). These results indicate that the benefits of not having parasites result in higher mating success under low-density conditions with little male-male interference competition and increased strength of selection against parasite load under such conditions.

**Table 1.**
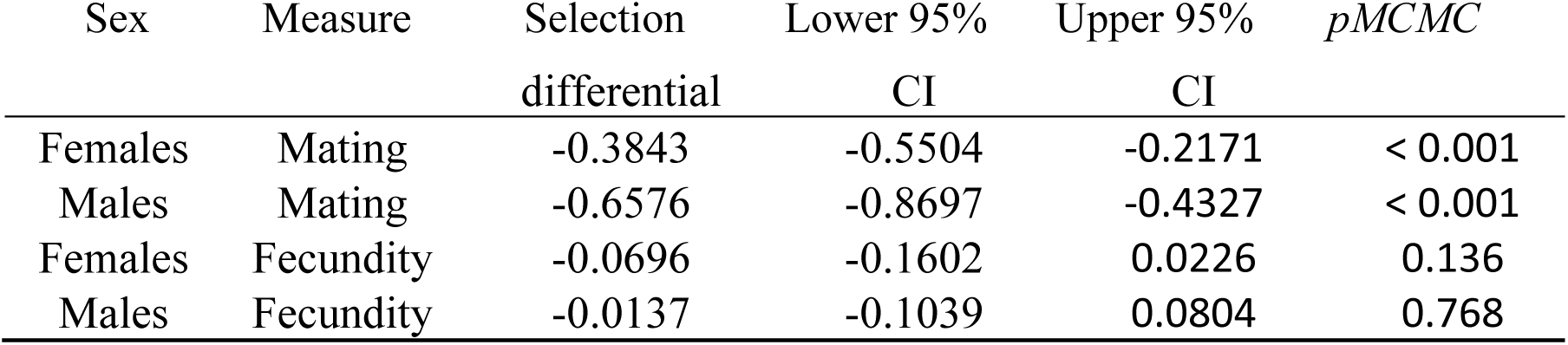
Precopulatory (mating success) and postcopulatory (fecundity) linear selection differentials, 95% confidence intervals and significance for parasitizedmales and females.

**Table 2.**
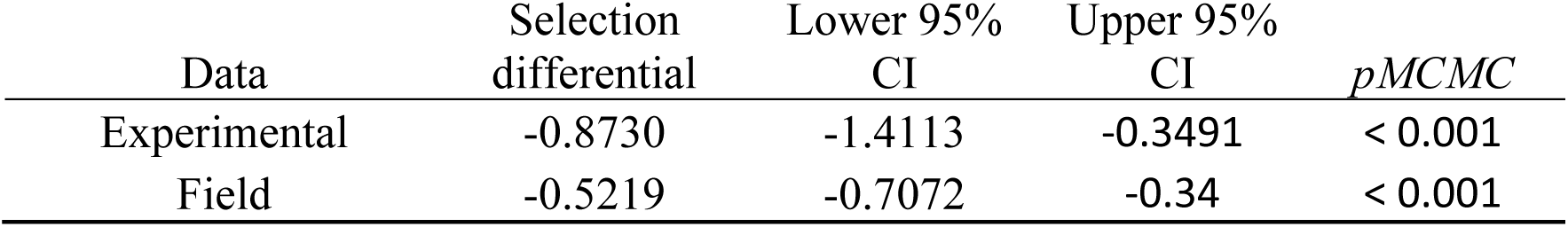
Precopulatory (mating success) linear selection differentials, 95% confidence intervals and significance for parasitized males in high-density field and low-density experimental conditions.

| Data | Selection differential | Lower 95% CI | Upper 95% CI | *p*MCMC |
| --- | --- | --- | --- | --- |
| Experimental | -0.8730 | -1.4113 | -0.3491 | < 0.001 |
| Field | -0.5219 | -0.7072 | -0.34 | < 0.001 |

### Population decline and temporal changes in parasite resistance and parasite tolerance

The population density in Lomma has declined significantly and steadily since 2003, as shown by the strong effect of year (pMCMC = 0.004; Fig 3, Supplementary table 3A). There was no sign of population recovery, as a quadratic term was not significant (results not shown). During the same time period (2003-2018), we found evidence for increased parasite resistance in both males and females as revealed by lower parasite numbers (Fig. 4A: pMCMC = 0.006). There was a significant sex difference, with females being more infected than males (pMCMC < 0.001), but no effect of the interaction between sex and year (pMCMC = 0.4; Fig. 4A. This indicates that parasite resistance has decreased in a similar magnitude in both sexes during these 15 years (Fig 4A, Supplementary table 3B).

**Figure 3.**
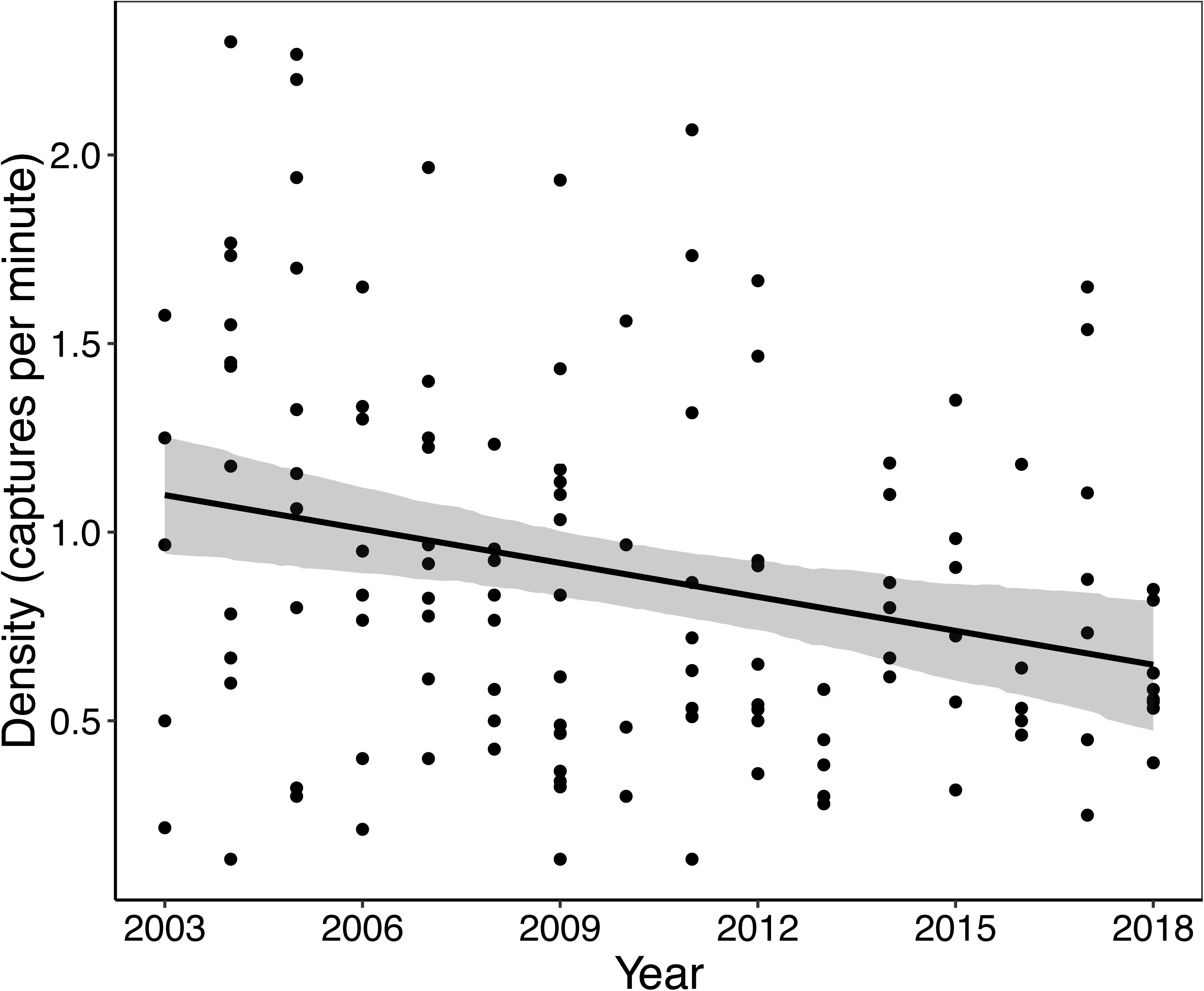
The population density of *I. elegans* in Lomma has decreased significantly since 2003. Points show population density in each visit, estimated by dividing the total catch (number of individuals) by the sampling effort (time spent catching) per visit. Regression line show predicted values from Bayesian models assuming a normal distribution, and shaded are 95% confidence intervals.

**Figure 4.**
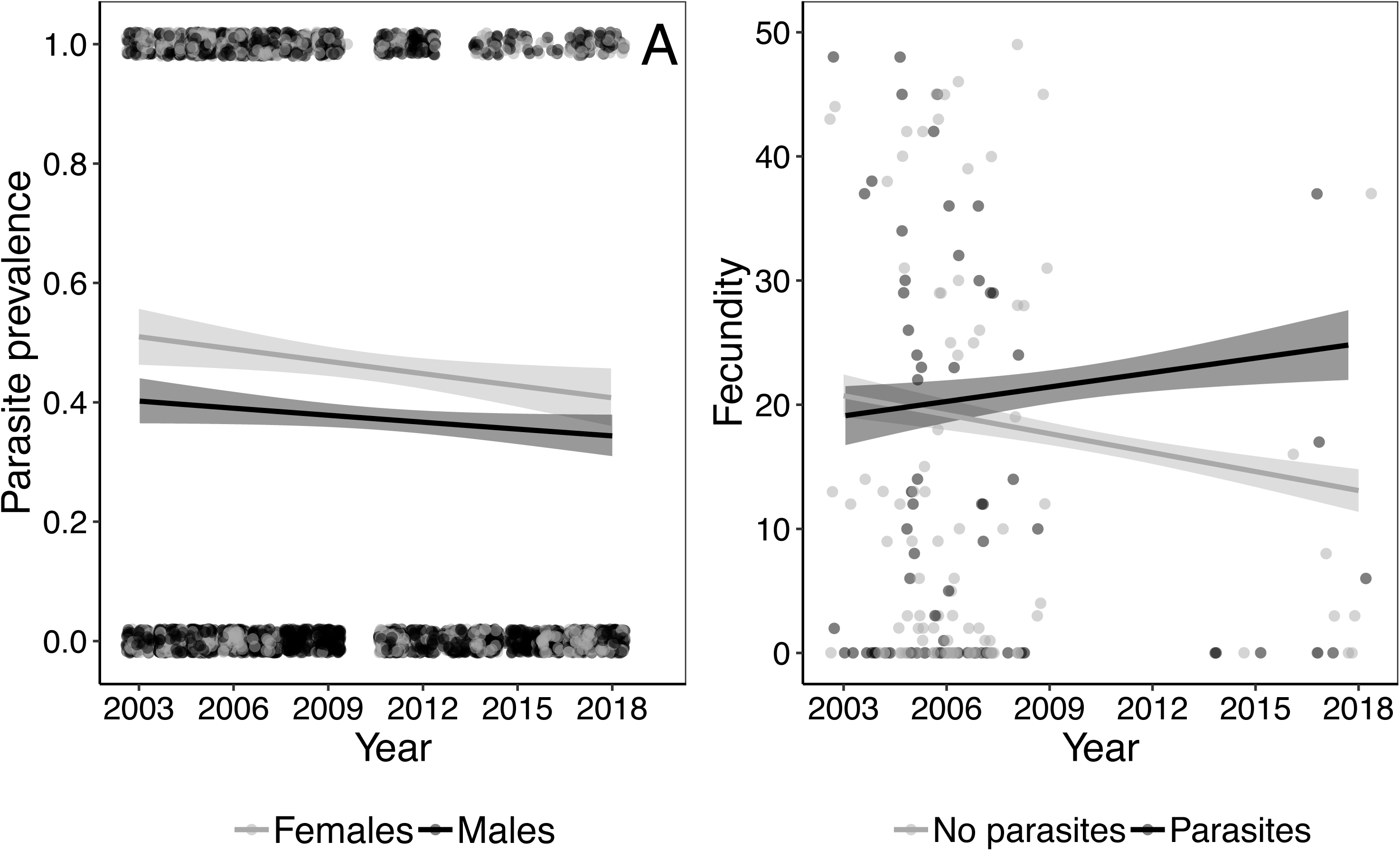
Temporal changes in parasite resistance (i.e. parasite prevalence) in both sexes **(A)** and male fecundity tolerance to parasites **(B).** Parasite prevalence has decreased over time, and this decline across the 15 years of study was similar for males and females **(A)**. Parasite tolerance, the reduction in the negative effect of parasite load, was analysed by coding males and females as “with parasites” or “no parasites”. We found a significant interaction of parasite prevalence in males and year, but no effect on females. The results show that parasitized males have increased their fecundity, with a steep positive effect in recent years **(B)**. We performed both models using a Bayesian framework assuming a binomial (for parasite prevalence) and Poisson (for parasite resistance) distribution. Points show individual observations and the regression line show predicted values with 95% confidence intervals in shaded areas.

We also we analysed temporal changes in parasite tolerance using our fecundity data. We found a significant effect of the interaction between year and parasite prevalence in males (pMCMC = 0.048). Fecundity of parasitized males increased during the time period, while non-parasitized males remain constant (Fig 4B). Interestingly, in females, there was no such temporal change and no significant change during the time period (pMCMC = 0.26, Supplementary table 3C).

## DISCUSSION

Theory and limited empirical evidence suggest that condition-dependent sexual selection can improve population fitness, for instance by purging deleterious mutations or by favouring locally adapted males [1,2,4,61,62]. Most previous studies investigating the links between sexual selection, local adaptation and population fitness have been performed in laboratory settings using a few model organisms and a restricted set of taxa [8–10,16,61–66].

Here, we used long-term population data on individual variation in parasite load as a measure of condition to investigate the role of sexual selection in promoting local adaptation of males and females, following the approach in some previous studies on parasite-mediated and condition-dependent sexual selection [37,38]. Overall organismal condition is influenced by both environmental and genetic factors, as condition is likely to reflect genetic variation across the entire genome [30]. Sexual selection on condition-dependent traits could therefore improve local adaptation, by purging novel deleterious mutations from the population [1,4]. Parasite-mediated sexual selection is therefore of general interest in this context, as individuals with low parasite load in a population could be argued to be locally adapted, having higher condition and presumably also being of higher genetic and/or phenotypic quality. For instance, sexual selection could reduce the risk of extinction by facilitating local adaptation if selection in males is stronger than in females, as the frequency of deleterious mutations in the population would decrease by purging selection on males, without reducing mean population fitness [4,10,17,18].

The results in this study show that sexual selection against parasitism, acting through mating success, is stronger in males than in females, while no effect of parasite load in postcopulatory selection through fecundity was found (Fig. 1B). This latter results contrast with the findings from a previous study where a significant negative effect of these ectoparasites on the fecundity of the different *I. elegans* female morphs was found [46]. The discrepancy between this previous study and the present one is partly due to the smaller sample size in the present work, where we used only data from complete male-female mating pairs from one relatively heavily infested population (Lomma). It is therefore possible that negative effects of ectoparasites on female fecundity would be only detectable in a larger dataset [46]. The proportion of parasitized females and males in the present study (47% and 38%, respectively) more than doubles the proportion of parasitized females and males in Willink and Svensson (21% and 14%, respectively [46]). The finding in the present study therefore reveals the effects of parasite load at a local scale in a population with high parasitic pressure, rather than the effects on a regional scale. The difference at local and regional scale as well as the temporal increase in parasite resistance and tolerance in this study (Fig. 4) strongly suggest ongoing local adaptation in the Lomma population as a response to high parasite pressure.

Parasite resistance (i.e., minimizing parasitic infection) and tolerance (i.e., reduction of the fitness costs of parasite load) are often viewed as alternative defence mechanisms in which investing in one will make the other defence mechanism redundant [67–69]. The results in the present study are consistent with such a trade-off in females, as parasite resistance has increased during the time period (Fig. 4A), whereas tolerance has not changed. In males, however, there is no evidence of such trade off, as both parasite resistance and tolerance have increased (Figs. 4A-B). Interestingly, there is yet no evidence that these changes in parasite tolerance and resistance have affected population growth, as population density has steadily declined (Fig. 3). This could potentially be explained by a temporal lag between increasing resistance and tolerance and their positive effects in population growth. If this is the case, we would expect population density to increase in the coming years. However, as a caveat, mean population fitness is likely to be more closely linked to female than to male fitness [16], so the effect of increased male tolerance on population size is by no means clear. Furthermore, population growth is also likely to be affected by several other biotic and abiotic factors, which could obscure the positive effects of increasing local adaptation to these parasites. Continued monitoring of this population in the future will hopefully enable us to better clarify the effects of ongoing local adaptation against parasites for population growth.

Condition-dependent sexual selection has been suggested to promote local adaptation, thereby reducing extinction risk in changing environments [11,15]. However, recent evidence from a laboratory experimental evolution study suggests that sexual selection cannot shield populations completely from extinction if environmental changes are too rapid [15]. The prospect for evolutionary rescue by sexual selection is also critically dependent on other factors, such as whether sexual selection is aligned (or not) with environmental change. If sexual or natural selection is frequency-dependent and aligned with the direction of abiotic environmental change, sexual selection can accelerate evolutionary rescue and thereby prevent population extinction [20].

Sexual selection could be important in evolutionary rescue, but the effects could be modulated by density-dependence as well as by other abiotic and biotic environmental factors. For instance, in low-density populations, genetic drift could increase the frequency of deleterious mutations reducing mean population fitness [8], an effect that could be counteracted if sexual selection reduce the mutation load and thereby promote evolutionary rescue. Empirical evidence of density-dependent sexual selection is contentious. At least one previous study found no evidence of density-dependent effects on sexual selection [8], whereas other studies found that sexual selection against males was stronger than in females in more complex environments with higher densities [9,70]. In a study of density-dependent sexual selection on horn length in fungus beetles (*Bolithotherus cornutus*), Conner [34] showed that sexual selection for long-horn males was stronger at low than at high density. A recent review of the limited available empirical evidence for how density influenced the strength of sexual selection revealed that that density could either increase or decrease the strength of selection, depending on natural history details and ecological conditions [33].

In this study, we found evidence for stronger sexual selection against parasite load in our experimental trials under low-density conditions, with fewer males, compared to the field conditions with more intense male-male scramble competition for females (Fig. 2). Male-male competition in our mating trials occurred exclusively between a non-parasitized (higher condition) male and a parasitized male (lower condition). The outcome of this experimental setup increased the relative mating success of non-parasitized males relative to field conditions, where, presumably, interference competition from other non-parasitized males are likely to reduce the relative fitness advantage of being non-parasitized [34]. Mating success therefore decreased more steeply with increasing parasite loads in this experimental low-density setting, compared to the higher density conditions in the field (Fig. 2A). Our results suggest that increased strength in sexual selection in males at low densities could facilitate evolutionary rescue. In this context, we note that the declining population size that we observed during the study period (Fig. 3) could increase the efficacy of parasite-mediated sexual selection on males (Fig. 2). By extension, such a link can lead to an eco-evolutionary feedback loop where an ecological factor (declining population density due to parasitism) could initially strengthen selection for local adaptation, subsequently increasing population density (cf. Fig. 4); and high population density would be expected to weaken parasite-mediated selection (cf. Fig. 2).

The beneficial effect of sexual selection promoting adaptation can be hindered by sexual conflict if male-induced harm reduces female fitness and the advantage of high-condition (presumably good genes) females [24,71,72]. Parasitism could potentially reduce male harm on females if parasitized males cannot pay the costs of searching and competing for females. Female fitness costs of male-induced harm would then be limited, and adaptation through sexual selection would be promoted. The potential for parasitism reducing male mating harassment and thereby reducing sexual conflict could be a fruitful new line of research.

In summary, our results show that parasite-mediated condition-dependent sexual selection is stronger in males than in females, and our longitudinal field population study was suggestive of ongoing local adaptation for increased parasite resistance in both sexes and increased parasite tolerance in males. We suggest that sexual selection against parasitized males becomes stronger under low-density than under high-density conditions, which opens up for eco-evolutionary feedback loops between density, parasite-mediated sexual selection and local adaptation. Low-density populations face a higher extinction risk, and condition-dependent sexual selection for locally adapted and non-parasitized males could potentially promote evolutionary rescue in this and other species. More generally, the links between parasitism, sexual selection, density, local adaptation and evolutionary rescue deserve further theoretical and empirical attention [1,3,14,20].

---

**Supplementary table 1.** MCMC binomial model of mating success.

|  | Posterior mean | Lower CI | Upper CI | *p*MCMC |
| --- | --- | --- | --- | --- |
| (Intercept) | -0.7818 | -1.738 | 0.0159 | 0.093 |
| Parasite load | -0.0307 | -0.0447 | -0.0175 | < 0.001 |
| Sex (males) | -0.6949 | -1.179 | -0.1957 | 0.0102 |
| Parasite load: Sex (males) | -0.0243 | -0.0476 | -0.0017 | 0.0310 |

**Supplementary Table 2.** MCMC model of fecundity

|  | Posterior mean | Lower CI | Upper CI | *p*MCMC |
| --- | --- | --- | --- | --- |
| (Intercept) | 2.9678 | 2.5459 | 3.3931 | < 0.001 |
| Female parasites | -0.0246 | -0.0647 | 0.0116 | 0.209 |
| Male parasites | 0.0093 | -0.0429 | 0.067 | 0.745 |

**Supplementary Table 3.** Binomial model of how male mating success is affected by parasite load under different settings (experimental low-density conditions and observational field data with high male-male scramble competition.

|  | mean |  |  |  |
| --- | --- | --- | --- | --- |
| (Intercept) | 1.9718 | 0.9553 | 3.0011 | < 0.001 |
| Parasite load | -0.1072 | -0.1512 | -0.0635 | < 0.001 |
| Data (observational) | -3.4744 | -4.2737 | -2.6969 | < 0.001 |
| Parasite load: Data (obs) | 0.0513 | 0.0026 | 0.0974 | 0.0254 |

**Supplementary Table 4.** Models of the change of population density (A), parasite resistance (B) and tolerance (C) through 15 years of population monitoring.

| **A)** | Posterior mean | Lower CI | Upper CI | pMCMC |
| --- | --- | --- | --- | --- |
| (Intercept) | 1.0982 | 0.9438 | 1.2558 | < 0.001 |
| Year | -0.0299 | -0.0489 | -0.0113 | 0.002 |

| **B)** | Posterior mean | Lower CI | Upper CI | pMCMC |
| --- | --- | --- | --- | --- |
| (Intercept) | 0.0487 | -0.1353 | 0.22 | 0.581 |
| Year | -0.0335 | -0.0584 | -0.0098 | 0.006 |
| Sex (males) | -0.5263 | -0.762 | -0.2908 | < 0.001 |
| Year: sex (males) | 0.0135 | -0.0187 | 0.0444 | 0.404 |

| **C)** | Posterior mean | Lower CI | Upper CI | pMCMC |
| --- | --- | --- | --- | --- |
| (Intercept) | 3.322 | 2.7134 | 3.9688 | < 0.001 |
| Year | -0.0157 | -0.1275 | 0.101 | 0.783 |
| Female (parasitized) | -0.5671 | -1.384 | 0.2655 | 0.174 |
| Male (parasitized) | -0.3977 | -1.2506 | 0.4776 | 0.367 |
| Year: female (par) | -0.0818 | -0.2415 | 0.0664 | 0.296 |
| Year: male (par) | 0.1609 | 0.0027 | 0.3152 | 0.044 |

Author contributions
MGL, EIS and AN designed the study. MGL and AN carried out the experiments. EIS and his former field assistants and PhD-students collected the observational long-term data. MGL conducted the statistical analyses and wrote the text with substantial input from EIS and AN.

## Acknowledgements

Observational data used in this study has been collected with the help of numerous PhD-students, field assistants and interns over the years. Sofie Nilén and Emily Scott helped collecting *I. elegans* individuals in the mating trial experiments. EIS was financially supported by research grants from Gyllenstiernska Krapperupsstiftelsen, Olle Engqvist Byggmästare Foundation (postdoctoral scholarship to MGL), Lunds Djurskyddsfond and the Swedish Research Council (VR; grant no. 2016-03356).

